# GnnDebugger: GNN based error correction in De Bruijn Graphs

**DOI:** 10.1101/2025.05.07.652713

**Authors:** Marijo Šimunović, Mile Šikić, Anton Bankevich

**Author notes:** Contributing authors.

## Abstract

**Motivation:** Modern sequencing technologies have enabled the reconstruction of complete mammalian genomes from telomere to telomere. However, scaling this achievement to thousands of species and population-level studies remains a challenge. Key bottlenecks include the low quality of the draft assemblies and the high coverage requirements. In particular, reconstructing complete and accurate sequences of both haplotypes in diploid genomes is especially difficult since the sequencing depth is not always sufficient to properly reconstruct diverged regions. Inspired by the success of neural networks in extracting patterns from the data on a massive scale, we introduce a method for correcting errors in De Bruijn Graphs using Graph Neural Networks.

**Results:** Our model provides a reliable classification of edges into correct and erroneous, especially for diploid genomes with coverage depth **35** and lower. We demonstrate that these predictions can guide the downstream read error correction algorithm and genome assembly, ultimately allowing for more accurate genome assembly.

**Availability and implementation:** Both *GnnDebugger* (https://github.com/m5imunovic/gnndebugger) and LJA (https://github.com/AntonBankevich/LJA/tree/gnndebugger) are available on GitHub. Datasets used for training and testing of ML model are available at Zenodo: https://doi.org/10.5281/zenodo.15073168. HG002 reference and reads are available at https://github.com/marbl/HG002. Primates references and reads are available at https://github.com/marbl/Primates.

## 1 Introduction

The goal of genome assembly algorithms is to generate high-quality genome sequences from error-prone read sets. The two fundamental approaches used for genome assembly are the Overlap-Layout-Consensus (OLC) and the De Bruijn Graph (DBG) based assembly. Initially designed for different types of data – OLC for Sanger sequencing and DBG for Illumina sequencing – these methods were successfully adapted to PacBio high-fidelity (HiFi) reads in modern assemblers [4, 6, 13, 22, 27, 34]. Both OLC and DBG approaches construct a graph representation of the sequencing data - *assembly graph*. Genome chromosomes correspond to paths in assembly graphs. We refer to the collection of these paths as *genome path*. The assembly problem amounts to finding the genome path in the assembly graph. Errors in reads impact genome assembly differently for OLC and DBG approaches. In overlap graphs, vertices represent reads and edges represent overlaps between read sequences. Errors in reads necessitate consideration of imperfect overlaps, increasing the amount of spurious links between reads originating from distant parts of the genome. Error correction is typically addressed in a *preliminary stage*, before the assembly graph is built [6, 22]. In DBGs, vertices represent *k*-mers and edges correspond to (*k* + 1)-mers derived from reads. The unbranching paths are collapsed into single edges, with lengths defined by the number of (*k* + 1)-mers. Sequencing errors generate incorrect (*k* + 1)-mers, adding erroneous vertices and edges. However, the collapsing process typically compresses erroneous (*k* + 1)-mers originating from the same read error into a single edge. Many successful genome assembly tools [3, 4, 13, 41] build an assembly graph from uncorrected reads and rely upon *graph-based correction*, that classifies edges as either correct (part of the genome path) or erroneous (arising from sequencing errors).

Regardless of the approach, most error correction relies on *ad-hoc* heuristics. A common strategy involves experts analyzing assembly graphs to identify typical data patterns and translating them into the particular genome assembly subroutine, such as detection and removal of tips and bulges [41]. This process is labor-intensive and hard to scale to a wide variety of different species or a wider range of coverage in sequencing experiments. The graph contains a large number of possible error patterns that cannot be identified using the manual effort of a genome assembly expert.

The most straightforward way to classify edges into correct and erroneous is to measure the read support of a given edge. Edges with average coverage by reads below a given threshold are declared erroneous. Unfortunately, this approach produces both false positive and false negative results. First, systematic errors in low-complexity regions – known artifacts of HiFi reads [38] – can lead to high support for erroneous edges. Second, low coverage and high error rates in such regions often result in insufficient support for correct edges (see Suplementary Data.1).

An additional challenge is presented by diploid genome sequencing with low coverage. Genome coverage is divided into two haplomes. As a result, many regions are statistically expected to lack sufficient data for reliable reconstruction. However, by combining data from both haplomes, in most cases, it is still possible to reconstruct the genome sequence. Unfortunately, this approach often fails for regions with high divergence between haplomes. A recent study [42] has shown that assemblies of highly divergent immunoglobulin loci are often incomplete (containing only one haplome sequence) or even incorrect. We argue that overlooking this issue would hinder many future genome assembly applications, particularly in population genomics. Reconstructing only a single haplome of a genome would favor haplotypes with simpler structure, potentially obscuring complex structural variations in the genome.

The genome coverage by HiFi reads is limited by the SMRT cell output and genome size. Analyzing mammalian genomes published by the VGP consortium [28], we established that around 57% had coverage below 35x, yielding less than 18x coverage per haplome. Recent advances in PacBio sequencing chemistry have significantly increased the yield of a single SMRT cell. However, this improvement is likely to popularize the barcoded sequencing of multiple samples within a single SMRT cell, rather than increasing coverage for individual genomes. Therefore, understanding how many samples can be multiplexed without significantly compromising assembly quality becomes an even more critical question.

Machine learning-based approaches have been successful in uncovering hidden patterns in the data and are capable of performing various tasks at the near-human level [14][35]. Recently, Graph Neural Networks (GNNs) [29] have been gaining popularity in domains with unstructured data. The application of GNNs is not always straightforward. They are known to suffer from over-smoothing, when predicted values depend solely on the local neighborhood of a node or edge, or over-squashing, when information needs to propagate along longer ranges[1]. Despite the drawbacks, these models have been successfully applied in various domains [7, 15, 18, 40]. Given the abundance of high-quality annotated data, ML can solve hard problems in the domain of biology [12]. The recent completion of high-quality assembly references [17, 23, 37] provides an opportunity to utilize ML methods in genome assembly.

*GnnDebugger* is a GNN-based approach for detecting errors - “bugs” – in assembly graphs. *GnnDebugger* combines generic features of edge lengths and coverage with topological information to infer the *multiplicity* - number of times the genome path visits a given edge - in DBGs. These predictions can be utilized to improve the detection and correction of errors in reads, a crucial condition for the success of assemblers that use PacBio HiFi long reads. We provide a method to create training data automatically, without the need for human labelers, and a model for predicting edge multiplicities. The GNN-based approach improves the performance of error detection in low-coverage setting where the heuristics struggle. We demonstrate the effectiveness of this method by integrating the outputs of the ML model into an existing DBG assembler, replacing the heuristic algorithm’s output, and consequently improving assembly results.

## 2 Related work

Machine learning methods have been utilized in bioinformatics to solve various problems, such as nanopore sequence basecalling [25], methylation detection [32] or protein structure prediction [12]. However, the adoption in the area of genome assembly has so far been very limited. For example, in [33], Conditional Random Fields have been used to predict arc multiplicities in DBGs using short reads. Similarly, in [39], graph coloring via neural network is used for haplotype phasing. Most of these methods are developed in isolation, without considering integration in full genome assembly pipelines.

Another hindrance to using ML in genome assembly is the lack of standardized practices. Each genome assembler has its particular way of building the graph and resolving errors that are rarely compatible with other assemblers. This lack of interoperability makes it harder to adopt the improvements from one assembler to the others and to define a standardized feature set on which ML techniques could be applied. In the domain of OLC-based assembly, the GNNome Framework [36] recently demonstrated the potential of GNNs to unravel overlap graphs and correctly identify genome paths. Data samples for training were generated using the existing assemblers. Large assembly graphs were partitioned into smaller graphs with varying cluster sizes, and boundary nodes were masked during the training. A bidirectional GNN with residual connections based on residual convolutional architecture [5] was used to predict most probable correct path through the graph. To the best of our knowledge, graph neural networks have not yet been used in DBG-based assemblers.

## 3 Methods

### 3.1 Problem description

We use the graph neural network model to predict the multiplicities of edges in the assembly graph. Although downstream analysis uses only binary classification: multiplicity is 0 (erroneous edge) or multiplicity is at least 1 (correct edge), we have observed that solving a more complex problem of multiplicity estimation results in a more accurate classification. This approach outperforms the alternative based on simple binary training criterion [31]. The binary classification can be used as a guide for graph-based error correction algorithms. However, classification errors may result in fragmented assembly or even assembly errors. Figure 1 illustrates how errors in edge classification can result in a fragmented assembly even when one of the haplomes is reconstructed. Given enough samples with low coverage, we can use neural networks to learn the error patterns in such *error hotspots*, avoiding the need to explicitly parameterize the decision algorithm. Consequently, improved edge classification reduces the number of errors and results in an improved assembly.

**Fig. 1:**
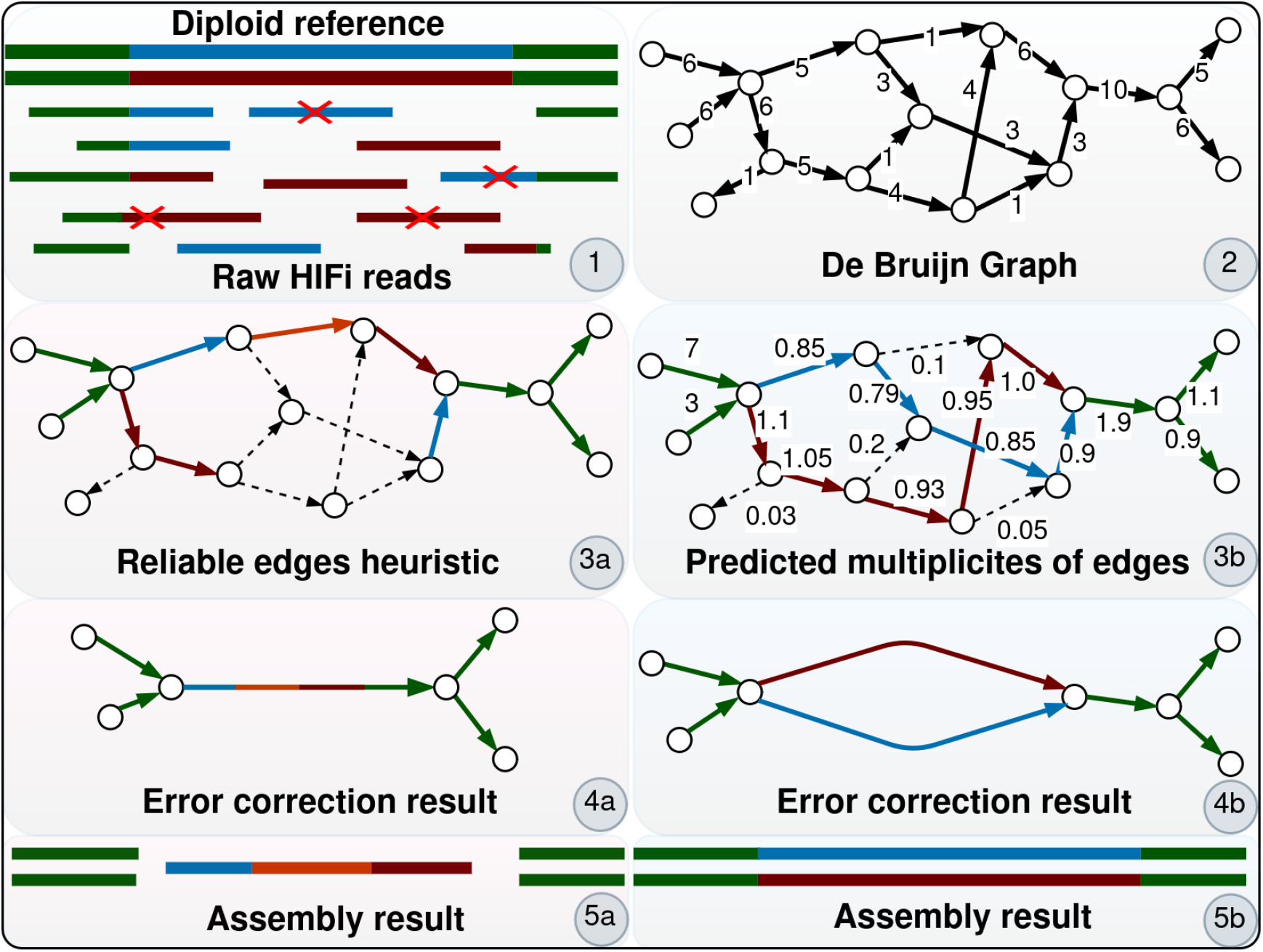
GnnDebugger predicts multiplicites of the edges in DBG graph and detects reliable edges. *1:* Diploid reference has been sampled with the ocassional errors in sequencing reads. *2:* DBG with edges formed by low coverage reads. *3a:* Error correction in heuristics misclassifies many edges due to low coverage. *3b:* The GNN is used to predict multiplicities in the graph. The predictions above threshold *t* = 0.5 are marked reliable (colored, full line) and erroneous (black, dashed line) otherwise. *4a:* Assembly process performed based on the *3a* input. *4b:*. Assembly process performed based on the *3b* input. *5a:* Lower accuracy of heuristics results in more errors and unresolved repeats. *5b:* Better error detection results in higher quality assembly.

### 3.2 Datasets

We use a highly accurate *HG002* reference to create datasets without resorting to laborious and expensive manual labeling. Each data sample is an assembly graph of a given unique set of input reads. We trained our model using two types of input reads: one, generated by simulating reads from a reference genome and the other, obtained by down-sampling a real read set. To simulate reads, we utilize the PbSim3 simulator [24]. First, we create simulation profiles from the sequencing reads of *CHM13* reference [23]. We simulate training and validation samples from different simulation profiles. In addition to simulated reads, we also utilize the HiFi reads which were used to assemble the *HG002* reference. Real reads are first mapped to respective chromosomes from references. We downsample the reads with different random seeds to generate graphs with varying coverage rates (see Table SI 5).

For our experiments, we always create a graph from a single diploid reference chromosome. For each assembly graph, we export edge index, edge features and multiplicity information. The edge features consist of edge coverage and edge length vectors. Coverage of an edge is the measure of its support by input reads. Formally, in de Bruijn graphs the coverage of a (*k* + 1)-mer is defined as the number of times it is present in input reads. Edge coverage is defined as the average coverage of all (*k* + 1)-mers in it.

Here, we describe our approach for labeling of the graph samples. Let *G* represent the assembly graph, *R* the reference genome, and *C* the homopolymer-compression function. For an edge *e* ∈ *G*, let *S*_*e*_ denote the sequence of edge *e*, and let *S*_*R*_ denote sequences (chromosome strands) within *R*. The homopolymer-compressed (HPC) sequence of *S*_*e*_ is *C*(*S*_*e*_), and homopolymer-compressed set of sequences in reference is *C*(*S*_*R*_). Then the label *L*(*e*) for edge *e*:

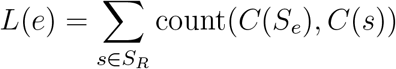

is the total count of occurrences of *C*(*S*_*e*_) in *C*(*S*_*R*_). Therefore, multiplicity is equal to zero if the edge does not appear in the reference, or a positive integer number if it appears one or more times in the reference. Multiplicity information is produced by matching HPC edges with the HPC reference. The main advantage of this labeling procedure is that it can be completely automated. One disadvantage is that occasionally, the edge can be labeled as false positive or false negative due to errors in the reference genome. Even highly curated benchmark datasets are known to have mislabeled data [20]. While errors in the test set can destabilize error benchmarks, our goal is not just to improve multiplicity prediction, but also the final assembly. Hence, we conducted additional evaluations with standard quality measures used in genomic assembly to evaluate the effectiveness of our method. Once the graph is assembled, we export the graph data in a format suitable for training and testing within the PyTorch framework [2, 8]. All datasets used in the article are available at [30].

#### 3.2.1 Data Transformations

The following transformations are applied to the data submitted to the model. For nodes, we add *in-degree* and *out-degree* counts. For edges, we normalize the length feature using *Z-Score* normalization. During the graph-based error correction, assemblers call the error correction routines iteratively. As the assembly graph goes through multiple corrections, the size of the edge lengths changes. Normalizing the length makes the model less sensitive to these variations. We also add two additional features, both boolean in nature: one indicating *single-* or *multiedge* attribute and the other, marking the edges at the *boundary* of the graph. We added the boundary edge feature due to significant coverage variability in those regions, where read support is typically lower. In addition to the node and edge features, we incorporate a global graph *coverage estimation* feature for the given input graph. GNN only processes small neighborhood of each respective edge, and cannot derive this information from local features. The typical coverage distribution in diploid assembly is bimodal. Most of the coverage values cluster around the first, higher peak, which corresponds to the coverage per haplome. The second peak is caused by homozygote regions of the two strands of the chromosome. We calculate a histogram of coverage values with 100 bins. Then, we detect the first peak, ignoring the bins with a coverage below 5 per haplome and use the 10 points surrounding the peak as a feature. Coverage less than 8 per haplome was not used in our generated datasets or in evaluation experiments.

### 3.3 Model

We use the model from [36] as an architectural basis and adapt input and output stages to the particularities of the DBG-based genome assembly approach. The model comprises an embedding stage, which converts input node and edge features into high-dimensional embedding vectors. A core part of the model is a GNN utilizing a message passing mechanism [9]. Unlike most existing GNN architectures, this model has been adapted to work bidirectionally, so that the predictions are made based on both upstream and downstream edge neighborhoods. Finally, a regressor stage predicts the edge multiplicity value. The regressor incorporates MLP-like element that takes the graph *coverage estimation* feature, converting it to a high-dimensional embedding. The graph feature aids the network in calibrating the multiplicity predictions to the variations in input read set coverage. Edge, node and graph features are concatenated into a single feature from which the final predicted value is calculated. Model implementation details are available in Suplementary Data.2

### 3.4 Objective

The main output of the graph-based error correction procedure is a boolean decision on whether the edge is reliable. Therefore, an equivalent approach would be to utilize an objective for binary classification, as in [36]. Initially, we reduced the ground truth multiplicity targets to binary choice by applying the boolean operator to the multiplicity value of the respective edges. However, collapsing multiplicity values results in a loss of valuable information that neural network can learn. Furthermore, multiplicity value predictions offer opportunities to improve the error correction and resolution of repeats in the graph [26]. Therefore, we decided to model our problem as a regression task. Our loss function combines Huber loss [11] and an additional regularization term, which we call *Flow factor*.

The loss function is defined as:

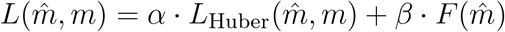

where *m* and *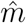* are ground truth and predicted multiplicity value, respectively. *α* and *β* are hyperparameters. *L*_*Huber*_ is Huber loss of the predicted edge multiplicity with parameter *δ* = 3.

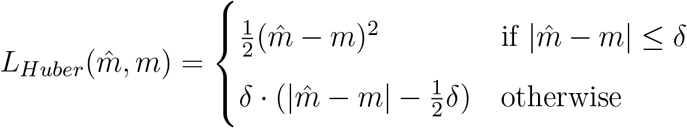

We use Huber loss in favor of MSE loss because the former is more sensitive to the miss-predictions of small multiplicities (up to the *δ*), which is really important as we set the decision boundary of whether the edge is correct or incorrect between 0 and 1. Using a linear loss term outside of the *δ* range reduces the sensitivity to outliers caused by repeats in the reference. Quadratic loss values would have a huge impact on the loss value without much benefit to the process of detecting erroneous edges. The idea of using flow is not new in DBG applications [26]. It is also used in [33] to train CRFs for multiplicity prediction. Additionally, it has been used for algorithmic reasoning based on GNNs [21]. Unlike the previous works, we use it with a little twist. It is not a loss term in a classical sense as we are not comparing the predicted values to a ground truth. It is a regularization term that favors preserving the flow in the graph. The flow loss of a single node *v* is defined as:

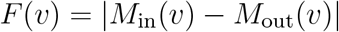

where *M*_*in*_(*v*) is a sum of multiplicities incoming from the neighboring nodes of node *v* and *M*_*out*_(*v*) is a sum of the multiplicities outgoing to the neighboring nodes of node *v* in directed graph G.

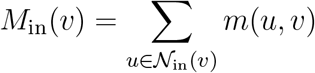

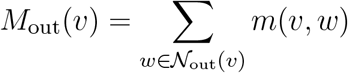

The total flow is then reduced by dividing it by the number of nodes *N* in the graph. The flow factor is important due to the way on how we create ground truth values for multiplicity. A small percentage of the edges, depending on the species, will have erroneous ground truth values. For example, homopolymer compression is going to make query of form “ACCCCTGTAT” hit the reference position with nucleotides “TACTGTATA”. Therefore, the misprediction will result in a lower penalty if the flow of the multiplicities in the local neighborhood of the node is preserved and presents a relaxation to the absolute prediction objective of the Huber loss. We recommend *α* = 0.3 and *β* = 0.7 as default settings based on the error analysis of the assemblies (see Table SI 1 and Table SI 2).

## 4 Results

### 4.1 Evaluation setup

#### 4.1.1 LJA genome assembly algorithm

The described methods could be applied to any DBG-based assembly algorithm. For the purposes of this research, the La Jolla Assembler (LJA)[4] was chosen based on the following considerations. First, LJA is currently the only genome assembly tool for HiFi reads that uses classic DBG instead of its sparse version. In this way, we avoid additional result variation caused by the choice of “anchor” k-mers [27]. Second, LJA does not use preliminary read correction, entirely relying on graph error correction, which removes the necessity of adjusting for possible bias and artifacts of preliminary error correction. Finally, the modular design of LJA allows the simple incorporation of alternative edge classification methods into its pipeline. *MowerDBG* module that is responsible for error correction in LJA uses algorithm 1 to correct errors in both the de Bruijn graph and reads. First, *mowerDBG* classifies edges into correct and erroneous based on their coverage and additional ad-hoc heuristics. Then, it attempts to reroute the paths of reads in the graph that go through erroneous edges. If all paths visiting erroneous edge are successfully rerouted, the edge is removed from the graph. Therefore, identifying reliable edges is the cornerstone of the *mowerDBG* algorithm.

##### Algorithm 1

mowerDBG Iteration(Graph, Reads)

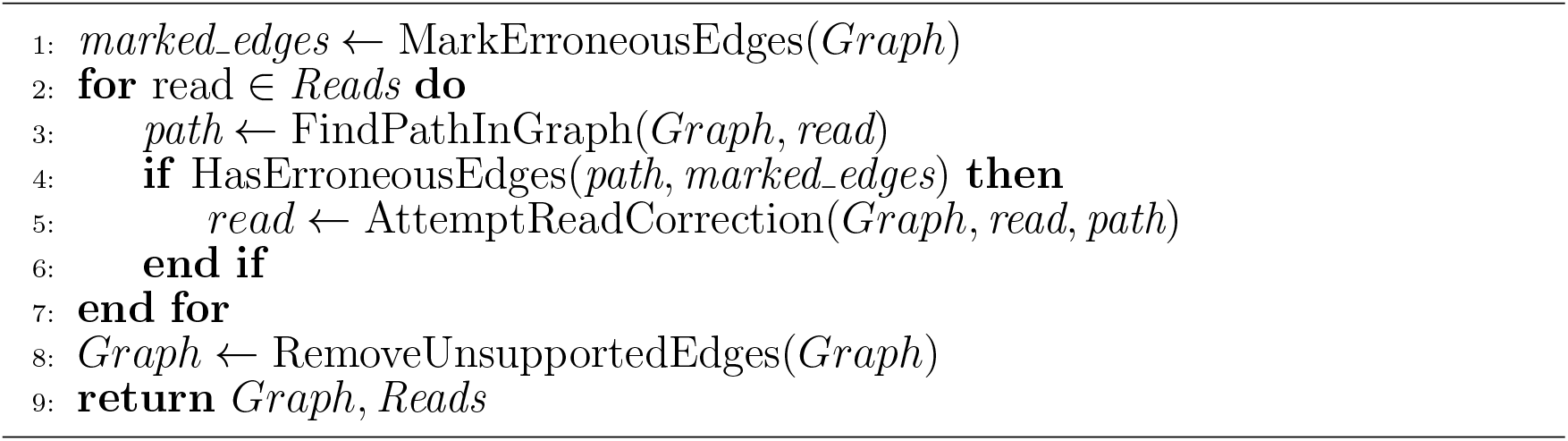

The algorithm 1 is applied several times during the error correction process, each with a different modification of step 5. In case there are multiple possible ways to correct a read, additional information (paths of other reads, edit distance, known patterns of errors in HiFi reads) may be used to make the choice. The parameters of this choice are changed from iteration to iteration to make the most reliable corrections first and to resolve more complex cases only after the majority of errors in the graph and in reads are already fixed. In total, this procedure is called 6 times during the assembly pipeline with 2 different values of *k* (501 and 5001).

To test the impact of the proposed edge classification on genome assembly results, we preserve the pipeline and parameters of the LJA error correction procedure but replace step 1 of each iteration with the criterion we developed using the GNN approach.

We evaluated assemblies based on assembly contiguity (NG50), number of contigs (LG50), rate of duplications in the assembly (Duplication ratio), structural errors (misassemblies), and mismatch/indel rate (mismatches and indels per 100Kbp). For precise descriptions of these metrics refer to [10]. Evaluation metrics were calculated using Quast-LG tool[19]. Unfortunately, no existing tool (including Quast-LG) can reliably detect structural errors in diploid genomes. Similarity between haplomes is confusing for assembly evaluation. We still report structural errors to emphasize the importance of assembly accuracy, but these values should be considered unreliable.

#### 4.1.2 Training

We trained models in our experiments for 300 epochs. The training data consists of 804 graphs in the training data (around 197 million edges), created from the chromosomes 4, 7, 8, 10, 11, 12, 17 and 20, respectively, and 334 graphs in the validation set, created from chromosomes 5, 16 and 19 (around 77 million edges), respectively (see SI 4, SI 5 for more details). Graphs were generated from read sets with coverage values from 8 to 17 per haplome. We used both uncorrected and semi-corrected reads (after 2 and 4 error corrections by *mowerDBG*). The network was trained using WAdam[16] optimizer with learning rate *lr* = 0.0005 in combination with *ReduceLROnPlateu* scheduler from the Pytorch library, which decreases the learning rate with *factor* = 0.95 if the validation error does not decrease for *patience* = 10 epochs. The GNN has 4 layers, with receptive field of 8 due to bidirectionality. Unless noted otherwise, internal hidden feature dimensions are set to 92. The same objective used in training was employed to measure validation set error for model selection.

#### 4.1.3 Evaluation stages

We first compared the performance of the *GnnDebugger* with the heuristic method implemented in LJA (see step 1 in Algorithm 1. For the evaluation of binary metrics (accuracy, precision, recall) we generated test datasets (see SI 3 for more details) from reads mapped to *HG002* chromosomes 3, 9, 18 and 21. We also tested our approach with two different *k* − *mer* settings: *k* = 501, and *k* = 5001. These values correspond to default parameters used in the LJA pipeline. The coverage of testing datasets ranges from 10 to 15 per haplome (20 to 30 for two haplomes). In addition, we generated a test set with coverage 21 per haplome for which the model was not trained. For each coverage, there were 14 graph samples tested. For the full assembly evaluation, we replace the heuristic method with *GnnDebugger*. The LJA constructs the assembly graph, exports it in the expected format[2, 8] and calls the inference procedure. The predicted multiplicities are read back into the LJA and assigned to the respective edges. For compatibility with the heuristic, we convert the multiplicity value to a boolean value based on the configurable threshold between 0 and 1 (see Suplementary Data.4 for recommended threshold settings). Based on these predictions, LJA partitions the paths in the graph and tries to reroute them to correct the errors in the reads.

For complete genome assembly, we used 3 diploid read sets from T2T-Primates project: *P. Troglodytes, P. Abelii* and *P. Pygmaeus*. The reads were downsampled to coverages 12, 15 and 21 per haplome, effectively creating nine different evaluation samples.

Finally, we conduct a series of model ablations to better understand the effect of various architectural decisions and data transformations,

### 4.2 Comparison with heuristic

The *GnnDebugger* significantly increases the accuracy and recall metrics while maintaining a similar performance on the precision metric (Figure 2). The heuristic outputs only binary values, so we performed the following conversion of the multiplicity predictions: Every prediction value higher than the threshold (*t* = 0.90) is converted to *true* value, i.e. we mark an edge as reliable and *false*, i.e. unreliable otherwise. The threshold is chosen to have a similar precision metric for *k* = 501, which makes it easier to compare other metrics, and is kept the same for *k* = 5001 datasets. In Figure 2, we can see that the median precision metric is similar for both the heuristic and the ML method, but the heuristic method has a much higher variance. The reason why heuristic has such high precision is a high cutoff threshold for coverage values. This means that there will be a very small number of false positive decisions, the trade-off being that many correct edges are false negatives. This has two drawbacks. One is that the error correction algorithms are pretty complicated in order to recover as much as possible of the reads information. The other is that the heterozygosity information often gets lost during the correction. We can demonstrate this by postprocessing all edges with average coverage less than 2.5 as false (green boxes in Figure 2). This results in further increase in precision at the cost of accuracy and recall loss. As the coverage decreases from 15 to 10, there is a significant drop of the accuracy - median accuracy drops 18% for the heuristic method, while the median accuracy of *GnnDebugger* only slightly (1.3%) decreases. The similar is the case for the recall. We can see that for the value of coverage 21, the metrics for heuristic equals the GNN model. In the case of the graph with *kmer* size 5001 we notice a similar pattern, with the difference that even for coverage 21 the *GnnDebugger* performs much better. The reason why the heuristic performs worse here is also connected to the properties of the graph. Longer *kmer* value means more unique edges with smaller coverage.

**Fig. 2:**
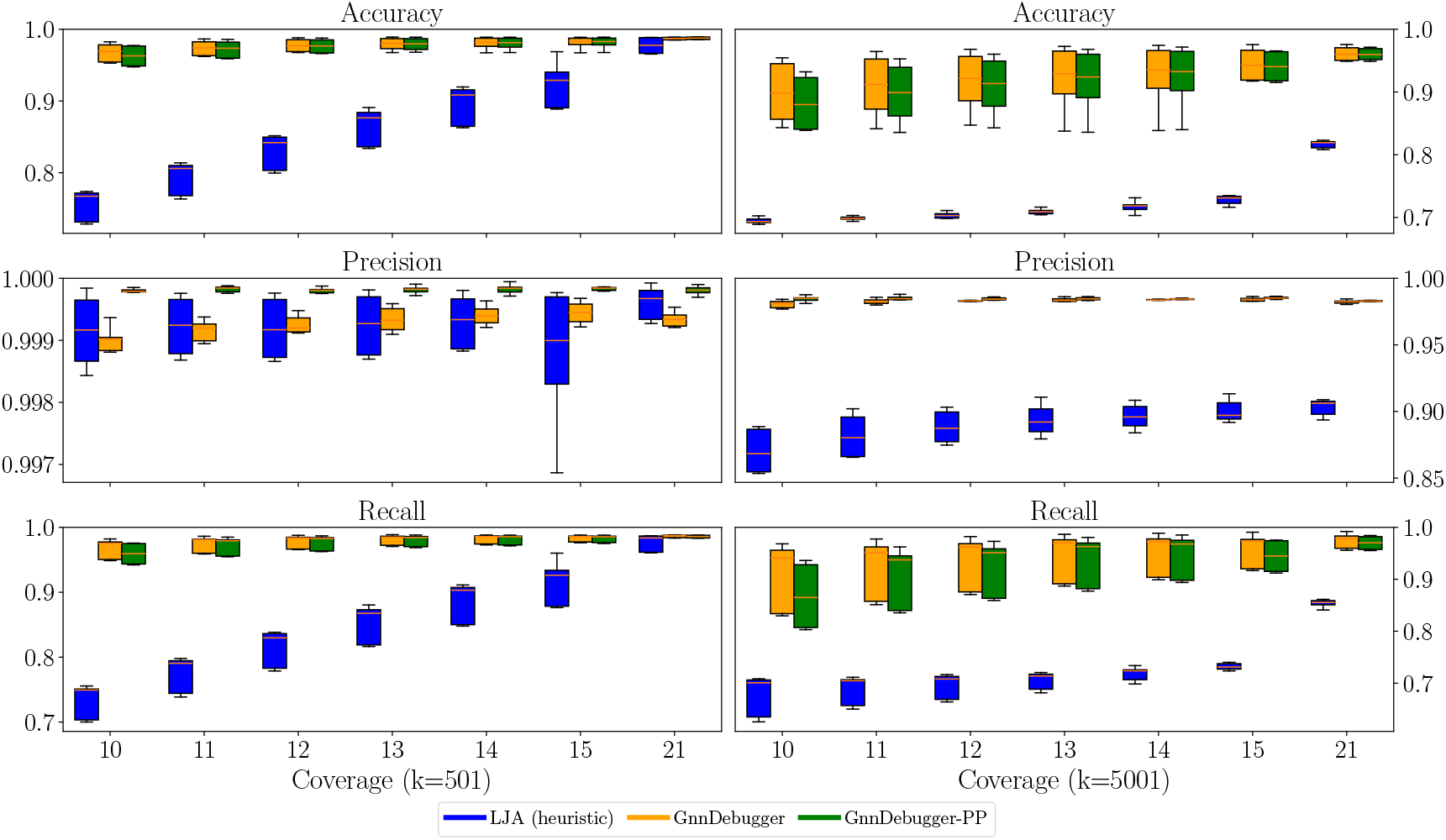
Accuracy, precision and recall metrics for DBG with *k* = 501 (left) and *k* = 5001 (right). Coverage values are given per haplome. The *LJA (heuristic)* error detection is threshold based. The *GnnDebugger* method uses a GNN network to classify the edges. *GnnDebugger-PP* marks all edges with average coverage below 2.5 as erroneous in a postprocessing step.

Most false positive mispredictions have small coverage values. Therefore, depending on the error rates of the assembler, we can easily further decrease the false positive rate by simply setting the edge reliability predictions in the range from 0 to 1 and small coverage to 0. However, performing such an operation has a detrimental effect, as there is substantial heterozygosity loss. In our experiments for full assembly, removal of false positives at the cost of removing true positives with small coverage had a negative effect on the error rates in assembly.

### 4.3 Evaluating results of complete assembly pipeline

We compared the results of LJA with the heuristic method (*LJA*) to the implementation relying on GNN detections (*LJA-GnnDebugger*). Additionally, we evaluated two other state-of-the-art assemblers, Hifiasm and Verkko, on the same read sets. Note that the Hifiasm does not output a completely compatible diploid assembly with those produced by LJA and Verkko. We used a haplotype-resolved processed unitig graph without small bubbles (*\*.bp.p utg.gfa* file), most similar to the output of the other two tools.

Figure 3 shows the metrics from the full assembly of *Pan Troglodytes, Pongo Pygmaeus* and *Pongo Abelii* references, respectively. In all experiments, we can see that the NG50 metrics and LG50 metrics are improved using the ML-based method. Hifiasm, and especially Verkko, have a bit higher duplication ratio in comparison to both versions of LJA assemblers. For coverage 15 per haplome, the number of misassemblies is comparable between *LJA-GnnDebugger* implementation and hifiasm. For lower coverage, the assembly quickly becomes very fragmented and unreliable. Verkko exhibited poor results on most parameters, which can be explained by Verkko focus on the hybrid assembly of PacBio HiFi and Oxford Nanopore reads. More detailed metrics are available in Suplementary Data.6.

**Fig. 3:**
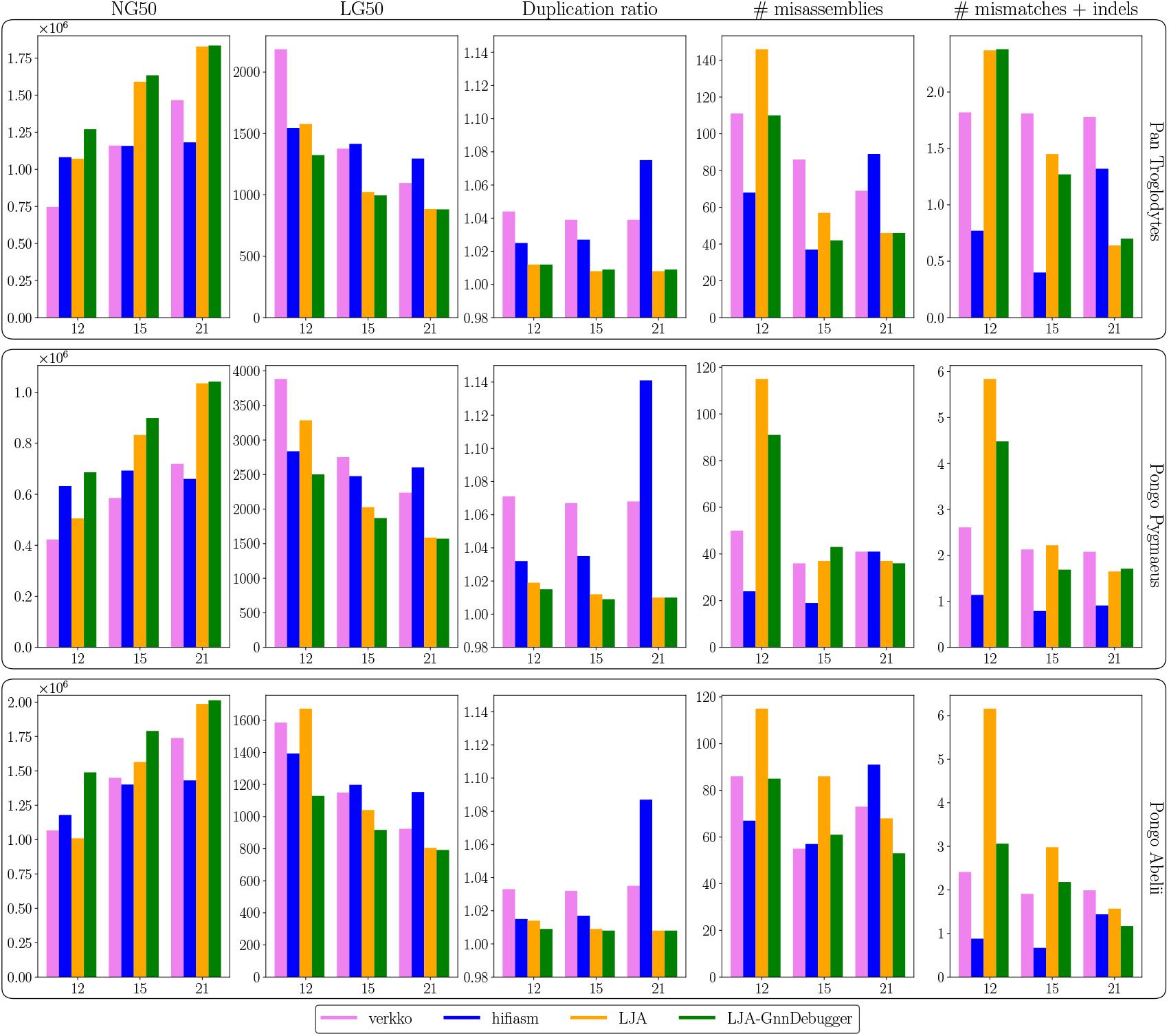
Evaluation of contiguity and accuracy of assemblies for *Pan Troglodytes (top), Pongo Pygmaeus (middle)* and *Pongo Abelii (bottom)* for coverages 12, 15 and 21 per haplome, respectively. Mismatches and indels are calculated per 100kbp.

*LJA-GnnDebugger* showed a significant advantage over LJA on almost all metrics and datasets. For assemblies with higher read coverage (21 per haplome) the LJA assembler with the ML based error correction improves assembly metrics over other assemblers in most of the evaluated cases. Therefore, the assembly quality significantly benefits from *GnnDebugger*.

### 4.4 Model ablations

The training was performed using the same dataset and the same optimization parameters as the baseline model, including the random seed. Figure 4 illustrates the performance of the models with various configurations. The removal of multiple layers leads to a general loss of performance. Notably, there was a significant drop in accuracy and recall metrics for the out-of-distribution samples when using a graph with *k*-mer size 501. Similarly, removing additional edge features (multiedge and tip) leads to a performance drop except for the recall metric in graphs with *k*-mer size of 5001. The overall effect of the global graph coverage attribute was challenging to estimate. For the graph with *k*-mer size 501 there was a slight improvement in accuracy and recall metrics, but precision was lower and exhibited much larger variation. Conversely, we observed a general improvement across all metrics for the graph with *k*-mer size of 5001. When the hidden feature size was decreased to 16, the model performance was similar to the baseline, suggesting that there is a redundant capacity to increase the size of the training dataset and incorporating additional reference species.

**Fig. 4:**
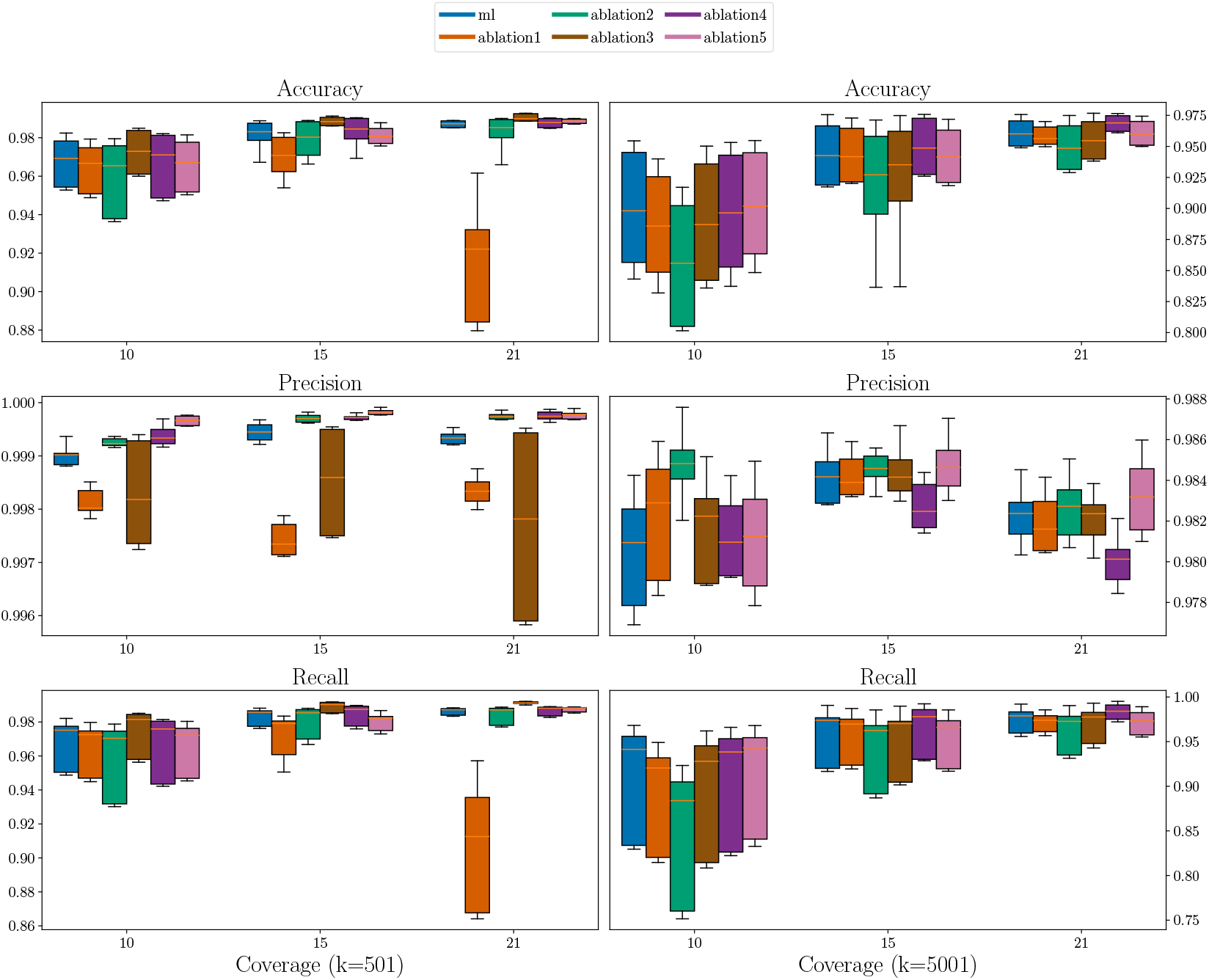
Model ablations for graphs built with *kmer* = 501 (left) and *kmer* = 5001 (right). The model without ablations is shown in blue. *Ablation* 1: The model is trained using a single GNN layer so that the network only sees the immediate *1-hop* neighborhood (orange). *Ablation* 2: removes edge features for multi-edge detection and tip detection(green). *Ablation* 3: removes the graph coverage estimation attribute (brown). *Ablation* 4: uses only 16 hidden features(purple). *Ablation* 5: does not use edge length scaling (pink).

## 5. Conclusion

ML based techniques have been able to improve the performance of many algorithms that have been relying on hand-crafted heuristics. There are, however, many hindrances to ML-adoption in genome assembly. An obvious problem is the lack of standardized data representations of genome assembly. Even though assembly graphs are a common denominator of most approaches, each genome assembler implements its own graph construction and error correction methods, making reusability and interoperability hard or even impossible. We tackled the problem of missing datasets for training and established a base architecture and learning objective for error correction in DBG based assemblers. We have shown how the ML-based approach can improve the detection and correction of errors in DBG-based assembly on low-coverage reads where traditional heuristic-based approaches face difficulties. Although our method uses very generic features, namely coverage and length of edges combined with topological information, it is still not guaranteed that other DBG assemblers will be able to provide this information for the ML algorithm.

An additional challenge arises from the limitations of genome assembly evaluation methods. The lack of tools to precisely detect structural errors in an assembly by comparing it to a reference raises significant concerns about the quality of mass-produced assembly results.

We hope to bring attention to these issues and spur a discussion in the direction of more theoretically grounded approaches in building assembly graphs and statistically based error correction methods, and establishing standardized interfaces and benchmark datasets for testing of each approach. Our future research will focus on improving the quality of the training data (mixtures of chromosomes (particularly acrocentric), various reference species, and regions of the genome known to be difficult to assemble). Our ablation studies showed that there is further research potential in identifying appropriate data features and model architectures for specific error detection problems. We also would like to try design error correction procedures that shall better utilize multiplicity predictions.

## Acknowledgments

This research was performed using the Advanced computing service provided by University of Zagreb University Computing Centre - SRCE, in scope of the project Geometrical Deep Learning for *de novo* Genome Assembly - NR-2024-01-010.

## 5.1 Author contributions

All authors contributed to research conceptualization, design of study, and manuscript preparation. *M.Sim.* prepared data, performed model training, testing, evaluation, and statistical analysis, and integrated the results into LJA. *A.B.* provided consultation on DBG-based genome assembly and LJA code, and *M.Sik* on machine learning. *M.Sik* and *A.B.* supervised the study. *M.Sik* provided computational resources.

## Competing interests

Authors declare no competing interests.

## Data and Code availability

*GnnDebugger* is available at https://github.com/m5imunovic/gnndebugger. LJA implementation with *GnnDebugger* can be obtained from https://github.com/AntonBankevich/LJA/tree/gnndebugger.

Training and test data for model production and evaluation is provided at Zenodo https://doi.org/10.5281/zenodo.15073168.

Primate references and reads are available from marbl/Primates. *Pan Troglodytes v2.0* reference is available at https://genomeark.s3.amazonaws.com/species/Pantroglodytes/mPanTro3/assemblycurated/mPanTro3.dip.cur.20231122.fasta.gz and HiFi reads at https://genomeark.s3.amazonaws.com/index.html?prefix=species/Pan_troglodytes/mPanTro3/assemblycurated/mapping/. *Pongo Pygmaeus v2.0* reference is available

HG002 reference and reads are available from marbl/HG002. The reference can be downloaded from HG002 v1.0.1 and reads from Revio HG002 v1.0.1. *Pongo Pygmaeus v2.0* reference is available at https://genomeark.s3.amazonaws.com/species/Pongo_pygmaeus/mPonPyg2/assemblycurated/mPonPyg2.dip.cur.20231122.fasta.gz and the HiFi reads from https://genomeark.s3.amazonaws.com/index.html?prefix=species/Pongopygmaeus/mPonPyg2/assemblycurated/mapping/. *Pongo Abelii v2.0* reference is available at https://genomeark.s3.amazonaws.com/species/Pongoabelii/mPonAbe1/assemblycurated/mPonAbe1.dip.cur.20231205.fasta.gzand the HiFi reads from https://genomeark.s3.amazonaws.com/index.html?prefix=species/Pongo_abelii/mPonAbe1/assemblycurated/mapping/.

The CHM13 HiFi sequencing data (32.4x, 20kbp libraries) is available at NCBI under accession numbers: SRX7897685, SRX7897686, SRX7897687, SRX7897688 and reference (v2.0) at GCF_009914755.1.

## Notes

### Competing Interest Statement

The authors have declared no competing interest.

