## Supplementary Data for "GnnDebugger: GNN based error correction in De Bruijn Graphs"

### Suplementary Data

#### Suplementary Data.1 Motivation for low coverage analysis in diploid genomes

In a genome sequencing project, there is an obvious objective of keeping coverage rates lower due to the lower sequencing costs. However, the problem of low coverage reads exists even in high-coverage datasets. Certain areas in a genome can form secondary structures that are difficult to denature and amplify [1]. T2T sequencing of the diploid human reference reports a coverage rate of 34.7 and most of the data within three standard deviations (7.03) [2]. This means that there is a substantial number of reads with low coverage. We determined that as much as 6.7 percent positions in chromosome 9 are covered less than 20 times. Lower coverage reads pose a specific challenge to DBG-based assemblers, particularly for diploid assemblies, resulting in collapses of heterozygous regions.

In cases where the sequencing is done with large coverage (e.g. at least 50), the heuristics approach provides accurate results. However, known recurring errors in HiFi reads sometimes lead to erroneous edges with significant coverage. For example, in the graph constructed from reads, originating from the HG002 chromosome 21, 1.55% erroneous edges have coverage larger than 3 when read coverage is 10 per haplome, up to 2.51% for read coverage 21 per haplome. Furthermore, graphs for diploid genomes frequently contain correct edges that have coverage 3 or less per haplome. For example, test samples from the HG002 chromosome 9 have 0.6% correct edges with coverage less than 3 when the sequencing depth is 21 per haplome up to 3.6% correct edges with coverage less than 3 when read coverage is 10 per haplome.

#### Suplementary Data.2 Machine learning model implementation

**Encoder:**

$\mathbf{W}_{n1}$, $\mathbf{W}_{n2}$, $\mathbf{W}_{e1}$ and $\mathbf{W}_{e2}$, $\mathbf{W}_{g1}$ and $\mathbf{W}_{g2}$ are learnable linear transformations. The same global graph feature is repeated for all edges.

$$\mathbf{h} = \mathbf{W}_{n2} \cdot \text{ReLU}(\mathbf{W}_{n1} \cdot \mathbf{x}_{node}) \tag{1}$$

$$\mathbf{e} = \mathbf{W}_{e2} \cdot \text{ReLU}(\mathbf{W}_{e1} \cdot \mathbf{x}_{edge}) \tag{2}$$

$$\mathbf{g} = \text{RepeatInterleave}(\mathbf{W}_{g2} \cdot \text{ReLU}(\mathbf{W}_{g1} \cdot \mathbf{x}_{graph}), edge\_index) \tag{3}$$

**Residual Gated Graph Neural Network:**

$\mathbf{A}_1$, $\mathbf{A}_2$, $\mathbf{A}_3$ and $\mathbf{B}_1$, $\mathbf{B}_2$ and $\mathbf{B}_3$ are learnable linear transformations. The node and edge embeddings $\mathbf{h}$ and $\mathbf{e}$ are passed into the graph neural network. Each GNN layer returns an updated embedding tuple ($\mathbf{h}$, $\mathbf{e}$), which is either passed into the next layer or encoder.

$$\mathbf{e} = \mathbf{e}_{j \to i} + \mathbf{e}_{i \to j} + \mathbf{e} \tag{4}$$

where

$$\mathbf{e}_{y \to x} = \text{ReLU}(\text{LayerNorm}(\mathbf{B}_1 \cdot \mathbf{e}_{y \to x} + \mathbf{B}_2 \cdot \mathbf{h}_y + \mathbf{B}_3 \cdot \mathbf{h}_x)) \tag{5}$$

$$\mathbf{h} = \text{ReLU}(\text{LayerNorm}((\mathbf{A_1} \cdot \mathbf{h} + \mathbf{h}_{fw} + \mathbf{h}_{bw})) + \mathbf{h} \tag{6}$$

For a single node $i$, forward and backward updates $\mathbf{h}_{fw}$ and $\mathbf{h}_{bw}$ are given by:

$$\mathbf{h}_{ifw} = \sum_{\mathbf{j} \in \mathcal{N}_{\mathbf{out}}(\mathbf{i})} \mathbf{A_2} \cdot \mathbf{h_i} \odot \sigma_{\mathbf{i} \to \mathbf{j}} \tag{7}$$

and

$$\mathbf{h}_{ibw} = \sum_{\mathbf{k} \in \mathcal{N}_{\mathbf{in}}(\mathbf{i})} \mathbf{A_3} \cdot \mathbf{h_i} \odot \sigma_{\mathbf{k} \to \mathbf{i}} \tag{8}$$

where $\odot$ is the Hadamard product. The gate $\sigma$ is calculated as the value of a sigmoid function of the edge feature of the incoming (or outgoing) edge divided by the values of the sigmoid functions of the edge feature for all incoming (or outgoing) edges.

**Decoder:**

The decoder concatenates the edge feature, the node features of an edge, and the global graph feature and calculates a dot product with a learnable vector W.

$$\mathbf{multiplicity} = \mathbf{W} \cdot [\mathbf{h}_{\text{src}}; \mathbf{h}_{\text{dst}}; \mathbf{e}; \mathbf{g}] + \mathbf{b} \tag{9}$$

#### Suplementary Data.3 Objective hyperparameters

We achieved the lowest error metrics with the models trained with $\alpha = 0.3$ and $\beta = 0.7$ as objective weights. Tables SI 1 and SI 2 contain the most important error metrics.

**Table SI 1**: Error metrics with respect to the coefficients of loss function weight parameters. The metric numbers are generated using Quast from assemblies coverage 15 per haplome

| | $\alpha = 0.1$ $\beta = 0.9$ | $\alpha = 0.2$ $\beta = 0.8$ | $\alpha = 0.3$ $\beta = 0.7$ | $\alpha = 0.4$ $\beta = 0.6$ | $\alpha = 0.5$ $\beta = 0.5$ | |
|---|---|---|---|---|---|---|
| misassemblies | 68 | 66 | 61 | 78 | 92 | *Pongo Abelii* |
| local misassemblies | 79 | 47 | 42 | 56 | 72 | |
| mismatches per 100 kbp | 2.09 | 2.41 | 2.18 | 3.38 | 4.81 | |
| indels per 100 kbp | 1.49 | 1.52 | 1.45 | 1.72 | 1.97 | |
| misassemblies | 34 | 40 | 43 | 49 | 79 | *P. Pygmaues* |
| local misassemblies | 97 | 106 | 76 | 137 | 148 | |
| mismatches per 100 kbp | 2.28 | 2.13 | 1.69 | 3.07 | 4.24 | |
| indels per 100 kbp | 2.23 | 2.23 | 2.05 | 2.46 | 2.72 | |
| misassemblies | 53 | 61 | 42 | 57 | 70 | *P. Troglodytes* |
| local misassemblies | 53 | 60 | 45 | 86 | 109 | |
| mismatches per 100 kbp | 1.26 | 1.25 | 1.27 | 1.84 | 2.09 | |
| indels per 100 kbp | 2.24 | 2.22 | 2.19 | 2.46 | 2.58 | |

**Table SI 2**: Error metrics with respect to the coefficients of loss function weight parameters. The metric numbers are generated using Quast from assemblies coverage 12 per haplome

| | $\alpha = 0.1$ $\beta = 0.9$ | $\alpha = 0.2$ $\beta = 0.8$ | $\alpha = 0.3$ $\beta = 0.7$ | $\alpha = 0.4$ $\beta = 0.6$ | $\alpha = 0.5$ $\beta = 0.5$ | |
|---|---|---|---|---|---|---|
| misassemblies | 94 | 87 | 85 | 119 | 147 | *P. Abelii* |
| local misassemblies | 135 | 122 | 104 | 158 | 163 | |
| mismatches per 100 kbp | 2.94 | 3.01 | 3.06 | 5.17 | 6.58 | |
| indels per 100 kbp | 2.71 | 2.73 | 2.66 | 3.15 | 3.40 | |
| misassemblies | 107 | 92 | 91 | 115 | 130 | *P. Pygmaeus* |
| local misassemblies | 297 | 265 | 245 | 337 | 399 | |
| mismatches per 100 kbp | 4.96 | 4.31 | 4.48 | 5.97 | 6.80 | |
| indels per 100 kbp | 4.94 | 4.72 | 4.57 | 5.28 | 5.59 | |
| misassemblies | 128 | 119 | 110 | 127 | 141 | *P. Troglodytes* |
| local misassemblies | 205 | 183 | 165 | 202 | 232 | |
| mismatches per 100 kbp | 2.16 | 2.37 | 2.38 | 2.81 | 3.67 | |
| indels per 100 kbp | 4.11 | 4.12 | 4.06 | 4.40 | 4.69 | |

#### Suplementary Data.4 Threshold selection

In order to determine the effect of the threshold setting, we run the assembly on the reads of HG002 chromosome 9 for multiple combinations of threshold settings in the first and second error correction stages. Our analysis shows that the model is relatively insensitive to the threshold settings in the second stage. However, in the first stage, setting the threshold to a value that is too high will increase the assembly error rates due to the collapse of the heterozygocity in the assembly. This also results in somewhat higher NG50 and lower LG50 values. Therefore, the values should be considered jointly when evaluating the results. We observe the same effect on full assembly metrics. We recommend a threshold of $t = 0.85$ for error correction stages with $k - mer = 501$ and $t = 0.70$ for error correction stages with $k - mer = 5001$.

Image SI 1 shows how metrics NG50, LG50, number of mismatches and indels change as we change the threshold value for first (small k-mer) and second (big k-mer) phase of error correction. Setting the threshold too high in the first phase typically results in a bigger error rate.

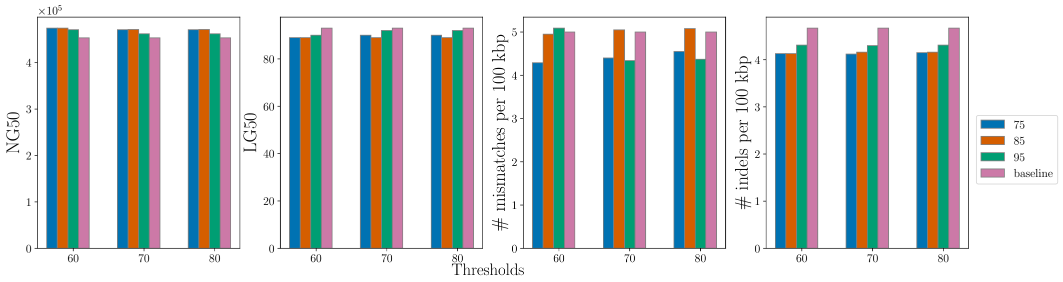


**Fig. SI 1**: Chromosome 9 Maternal strand. Metrics variability with varying thresholds for first (legend) and second error detection stage (x-axis).

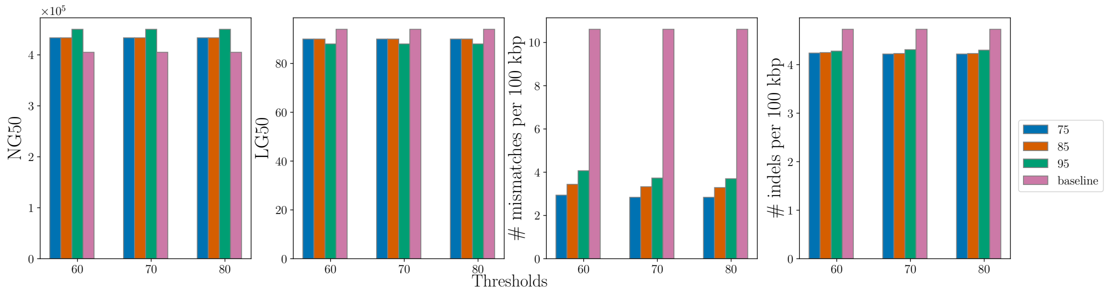


**Fig. SI 2**: Chromosome 9 Paternal strand. Metrics variability with varying thresholds for first (legend) and second error detection stage (x-axis).

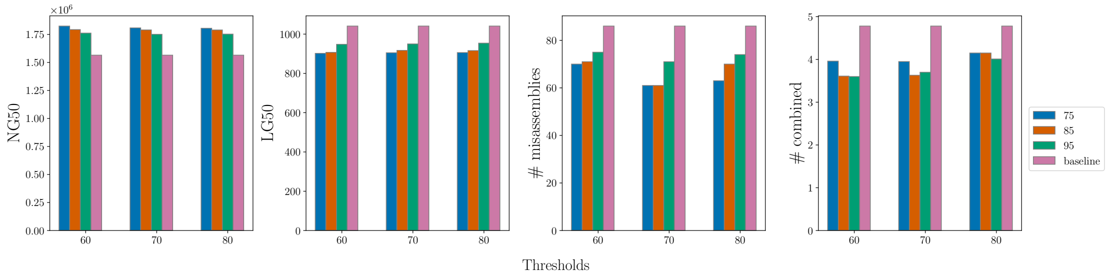


**Fig. SI 3**: Pongo Abelii. Metrics variability with varying thresholds for first (legend) and second error detection stage (x-axis).

#### Suplementary Data.5 Dataset summary

**Table SI 3**: Summary of test datasets. Each coverage contains 14 graphs created from reads mapped to chromosomes 3,9,18 and 21, respectively. Each edge in a graph is one evaluation point.

| | k=501 | | k=5001 | |
|---|---|---|---|---|
| Coverage | Nodes | Edges | Nodes | Edges |
| 10 | 2602886 | 3818236 | 221156 | 277880 |
| 11 | 2613490 | 3828870 | 209754 | 268034 |
| 12 | 2644708 | 3869347 | 201936 | 261354 |
| 13 | 2678022 | 3911718 | 196938 | 257868 |
| 14 | 2718148 | 3964823 | 195014 | 256804 |
| 15 | 2762886 | 4024712 | 192044 | 255159 |
| 21 | 3087462 | 4465576 | 197884 | 264706 |

**Table SI 4**: Summary of simulated train and validation graph statistics. Column *Chromosomes* contains information about which chromosome was used for simulating the graph and how many times it was used at specific coverage. Each edge is a training/validation point. Graphs are created with jumboDBG module of LJA assembler with k-mer size 501

| Samples | Nodes | Edges | Chromosomes (Samples) | K |
|---|---|---|---|---|
| 127 (train) | 35817438 | 52158422 | 4(16),7(16),8(15),10(16),11(16),12(16),17(16),20(16) | 501 |
| 48 (val) | 13040524 | 19029263 | 5(16),16(16),19(16) | 501 |

**Table SI 5**: Summary of downsampled train and validation datasets. Graphs are generated from raw reads and error corrected reads after two error correction iterations for k-mer size 501 and before the first of two error correction iterations for k-mer size 5001

| Samples | Nodes | Edges | Chromosomes(Samples) | Stage | K |
|---|---|---|---|---|---|
| 110 (train) | 23231274 | 33821746 | 4(16),7(15),8(16),11(15),12(16),17(16),20(16) | 0 | 501 |
| 36 (val) | 7192776 | 10498436 | 5(12),16(12),19(12) | 0 | 501 |
| Samples | Nodes | Edges | Chromosomes (samples) | Stage | K |
| 456 (train) | 72961560 | 109121336 | 4(66),7(65),8(66),11(66),12(64),17(65),20(64) | 2 | 501 |
| 202 (val) | 31302494 | 46847004 | 5(68),16(67),19(67) | 2 | 501 |
| Samples | Nodes | Edges | Chromosomes (Samples) | Stage | K |
| 111 (train) | 1564156 | 2180572 | 4(16),7(15),8(16),11(16),12(16),17(16),20(16) | 0 | 5001 |
| 48 (val) | 640054 | 879744 | 5(16),16(16),19(16) | 0 | 5001 |

**Suplementary Data.6 Assembly metrics for primate genomes**

**Table SI 6**: Pan Troglodytes assembly stats (reference length=6176211936)

| | Coverage 12 | | | | Coverage 15 | | | | Coverage 21 | | | |
|---|---|---|---|---|---|---|---|---|---|---|---|---|
| Assembly | LJA | hifiasm | LJA-GNN | verkko | LJA | hifiasm | LJA-GNN | verkko | LJA | hifiasm | LJA-GNN | verkko |
| Genome fraction (%) | 95.775 | 95.968 | 95.805 | 95.460 | 96.376 | 96.570 | 96.370 | 95.917 | 96.574 | 97.800 | 96.485 | 96.249 |
| GC (%) | 40.65 | 40.63 | 40.65 | 40.71 | 40.65 | 40.62 | 40.64 | 40.71 | 40.65 | 40.71 | 40.65 | 40.72 |
| Reference GC (%) | 40.60 | 40.60 | 40.60 | 40.60 | 40.60 | 40.60 | 40.60 | 40.60 | 40.60 | 40.60 | 40.60 | 40.60 |
| Duplication ratio | 1.012 | 1.025 | 1.012 | 1.044 | 1.008 | 1.027 | 1.009 | 1.039 | 1.008 | 1.075 | 1.009 | 1.039 |
| Total aligned length | 5974715762 | 6067165553 | 5979249338 | 6144800774 | 5993108351 | 6117102585 | 5993601663 | 6144319186 | 6005420301 | 6485069816 | 6003265117 | 6164938075 |
| NG50 | 1071091 | 1081648 | 1270206 | 746632 | 1590195 | 1157738 | 1633711 | 1159922 | 1827841 | 1181486 | 1834012 | 1466109 |
| NG90 | 121843 | 173037 | 136698 | 97143 | 165304 | 192832 | 177056 | 126804 | 196645 | 199972 | 195222 | 153059 |
| LG50 | 1578 | 1546 | 1324 | 2187 | 1024 | 1416 | 996 | 1377 | 885 | 1296 | 882 | 1097 |
| LG90 | 7215 | 6481 | 6095 | 10111 | 4860 | 6020 | 4661 | 6755 | 4158 | 5834 | 4163 | 5340 |
| # local misassemblies | 110 | 17 | 165 | 46 | 56 | 19 | 45 | 26 | 24 | 26 | 18 | 27 |
| # misassemblies | 146 | 68 | 110 | 111 | 57 | 37 | 42 | 86 | 46 | 89 | 46 | 69 |
| # mismatches per 100 kbp | 2.37 | 0.77 | 2.38 | 1.82 | 1.45 | 0.40 | 1.27 | 1.81 | 0.64 | 1.32 | 0.70 | 1.78 |
| # indels per 100 kbp | 3.95 | 2.00 | 4.06 | 7.67 | 2.26 | 1.19 | 2.19 | 5.89 | 1.56 | 1.52 | 1.62 | 5.15 |

**Table SI 7**: Pongo Pygmaeus assembly stats (reference length=6228071324)

| | Coverage 12 | | | | Coverage 15 | | | | Coverage 21 | | | |
|---|---|---|---|---|---|---|---|---|---|---|---|---|
| Assembly | LJA | hifiasm | LJA-GNN | verkko | LJA | hifiasm | LJA-GNN | verkko | LJA | hifiasm | LJA-GNN | verkko |
| enome fraction (%) | 94.963 | 94.934 | 95.570 | 94.551 | 96.235 | 95.320 | 96.470 | 95.263 | 96.610 | 97.537 | 96.576 | 95.798 |
| GC (%) | 40.91 | 40.91 | 40.91 | 40.93 | 40.92 | 40.92 | 40.92 | 40.95 | 40.93 | 40.92 | 40.93 | 40.96 |
| Reference GC (%) | 40.91 | 40.91 | 40.91 | 40.91 | 40.91 | 40.91 | 40.91 | 40.91 | 40.91 | 40.91 | 40.91 | 40.91 |
| Duplication ratio | 1.019 | 1.032 | 1.015 | 1.071 | 1.012 | 1.035 | 1.009 | 1.067 | 1.010 | 1.141 | 1.010 | 1.068 |
| Total aligned length | 6010252834 | 6082240721 | 6023314442 | 6292334741 | 6046536422 | 6129189740 | 6048466098 | 6316131599 | 6058513325 | 6913580987 | 6058059413 | 6353585168 |
| NG50 | 505488 | 632316 | 686297 | 422691 | 832476 | 692934 | 899075 | 585467 | 1035128 | 660315 | 1041919 | 719444 |
| NG90 | 76689 | 112637 | 88744 | 53343 | 110625 | 119298 | 116865 | 62185 | 141903 | 117751 | 141977 | 75482 |
| LG50 | 3285 | 2837 | 2502 | 3884 | 2028 | 2477 | 1870 | 2752 | 1587 | 2604 | 1572 | 2238 |
| LG90 | 14991 | 11306 | 11329 | 19696 | 8991 | 10302 | 8319 | 14621 | 6971 | 10674 | 6935 | 11460 |
| # local misassemblies | 188 | 9 | 245 | 21 | 58 | 11 | 76 | 16 | 37 | 26 | 32 | 25 |
| # misassemblies | 115 | 24 | 91 | 50 | 37 | 19 | 43 | 36 | 37 | 41 | 36 | 41 |
| # mismatches per 100 kbp | 5.84 | 1.14 | 4.48 | 2.61 | 2.22 | 0.79 | 1.69 | 2.13 | 1.65 | 0.91 | 1.71 | 2.08 |
| # indels per 100 kbp | 5.89 | 2.65 | 4.57 | 12.71 | 2.37 | 1.56 | 2.05 | 9.81 | 1.33 | 1.27 | 1.33 | 8.15 |

**Table SI 8**: Pongo Abelii assembly stats (reference length=6260850238)

| | Coverage 12 | | | | Coverage 15 | | | | Coverage 21 | | | |
|---|---|---|---|---|---|---|---|---|---|---|---|---|
| Assembly | LJA | hifiasm | LJA-GNN | verkko | LJA | hifiasm | LJA-GNN | verkko | LJA | hifiasm | LJA-GNN | verkko |
| Genome fraction (%) | 92.480 | 92.084 | 93.060 | 92.967 | 93.364 | 92.431 | 93.410 | 93.258 | 93.558 | 94.707 | 93.637 | 93.559 |
| GC (%) | 40.91 | 40.91 | 40.92 | 40.95 | 40.93 | 40.91 | 40.93 | 40.97 | 40.94 | 40.98 | 40.94 | 40.99 |
| Reference GC (%) | 40.91 | 40.91 | 40.91 | 40.91 | 40.91 | 40.91 | 40.91 | 40.91 | 40.91 | 40.91 | 40.91 | 40.91 |
| Duplication ratio | 1.014 | 1.015 | 1.009 | 1.033 | 1.009 | 1.017 | 1.008 | 1.032 | 1.008 | 1.087 | 1.008 | 1.035 |
| Total aligned length | 5857197683 | 5838436045 | 5864690300 | 5996292269 | 5881622997 | 5868307067 | 5882097538 | 6013342415 | 5891423172 | 6431946360 | 5891889759 | 6046900876 |
| NG50 | 1009253 | 1178749 | 1488691 | 1066772 | 1564865 | 1400258 | 1790037 | 1449275 | 1986125 | 1430299 | 2013578 | 1738702 |
| NG90 | 75277 | 85939 | 83173 | 78200 | 87282 | 92075 | 88396 | 87342 | 94916 | 95146 | 94732 | 98566 |
| LG50 | 1673 | 1393 | 1129 | 1586 | 1041 | 1198 | 917 | 1150 | 805 | 1153 | 792 | 924 |
| LG90 | 9123 | 7639 | 6432 | 8668 | 6042 | 6674 | 5528 | 6726 | 4999 | 6641 | 4949 | 5668 |
| # local misassemblies | 150 | 28 | 104 | 52 | 93 | 31 | 68 | 38 | 28 | 47 | 26 | 35 |
| # misassemblies | 115 | 67 | 85 | 86 | 86 | 57 | 61 | 55 | 68 | 91 | 53 | 73 |
| # mismatches per 100 kbp | 6.16 | 0.88 | 3.06 | 2.41 | 2.98 | 0.67 | 2.18 | 1.91 | 1.57 | 1.44 | 1.17 | 1.99 |
| # indels per 100 kbp | 4.28 | 1.59 | 2.66 | 5.23 | 1.80 | 0.90 | 1.45 | 4.37 | 1.02 | 0.90 | 0.93 | 3.80 |

#### Suplementary Data.7 Software commands

**Quast**: To calculate the assembly statistics we used quast (v5.2.0):

Listing 1: Quast invocation

```
#!/bin/bash
python3 quast.py \
        --large \
        --x-for-Nx 90 \
        --threads 32 \
        --debug \
        --skip-unaligned-mis-contigs \
        -e \
        --min-identity 98. \
        --min-alignment 10000 \
        --scaffold-gap-max-size 5000000 \
        --no-snps \
        --min-contig 15000 \
        <assembly.fasta> \
        -r <reference.fasta> \
        -l <label> \
        -o <out_dir> --no-plots --no-sv
```

**Hifiasm**: To generate assemblies using hifiasm (v0.20.0-r639) we run the following command:

Listing 2: Hifiasm assembly

```
hifiasm -t <threads> <reads.fastq>
awk "/^S/{print ">"$2;print $3}" \
hifiasm.asm.bp.p_utg.gfa > assembly.fasta
```

**Verkko**: To generate assemblies using verkko (v2.2-0) we run the following command:

Listing 3: Verkko assembly

```
verkko --threads <threads> --hifi <reads.fastq> -d <out>
```

**La Jolla**: To generate assemblies for LJA we used the command:

Listing 4: LJA assembly

```
# LJA (heuristic) assembly
lja --diploid -reads <reads.fastq> \
    -o <out> --threads <treads>
# LJA-GNN assembly
lja --diploid -reads <reads.fastq>  \
    -o <out> --threads <treads> \
    --CoverageBasedCorrection.reliability-mode multiplicity \
    --CoverageBasedCorrection.ml-threshold <t1> \
    --TopologyBasedCorrection.reliability-mode multiplicity \
    --TopologyBasedCorrection.ml-threshold <t2>
```

**seqtk**: For downsampling of the reads to desired coverage we use the seqtk tool (v1.3-r106) with following command:

Listing 5: Downsampling reads

```
seqtk sample -s <seed> <read.fastq> <fraction> > <out.fastq>
```

**samtools**: To extract the mapped reads from the respective *bam* file we used samtools (v1.13):

Listing 6: Extracting the reads from BAM file

```
samtools view -bh <reads.bam> <strand_id> -o <out.bam>
samtools fastq <out.bam> > <out.fastq>
```

**minigraph**: We used minigraph to map the contigs to chromosome strands in combination with *awk* and *seqtk*:

Listing 7: Mapping contigs to strands

```
minigraph -t <threads> -x asm <ref_chromosome.fasta> \
    <asm.fasta> > <graph.gaf>
awk -v var=<chr_id_hap1> '$6 == var {print $1}' <graph.gaf> \
    | sort -u > <hap1_contigs.txt>
awk -v var=<chr_id_hap2> '$6 == var {print $1}' <graph.gaf> \
    | sort -u > "<hap2_contigs.txt>
seqtk subseq <asm.fasta> <hap1_contigs.txt> > <hap1.fasta>
seqtk subseq <asm.fasta> <hap2_contigs.txt> > <hap2.fasta>
```

**coverage**: To estimate the coverage of the positions in reference we run the following script:

```
minimap2 -t <threads> -a -x map-hifi <ref.fasta> \
    <reads.fastq> > <alignment.sam>
samtools view -Sb <alignment.sam> <alignment.bam>
samtools sort <alignment.bam> -o <alignment.sorted.bam>
samtools index <alignment.sorted.bam>
samtools depth -a <alignment.sorted.bam> <cov.txt>
awk '{sum+=$3} END {print "Average coverage:", sum/NR}' \
    <cov.txt> > <avg_cov.txt>
```

##### Suplementary Data.8 Training and Inference

The following setup makes the assumption that the *GnnDebugger* and *LJA* repository is cloned to directory */work*, datasets are stored under */data* directory and configurations under */data/tconfig* and */data/iconfig* for training and inference configs, respectively. Visit GitHub pages of project for more detailed instructions.

```
git clone https://github.com/m5imunovic/gnndebugger
git clone https://github.com/AntonBankevich/LJA/tree/gnndebugger
# For training with GnnDebugger
cd /work/gnndebugger/apptainer
bash build_apptainer.sh
# For building LJA container with GnnDebugger
cd /work/LJA/apptainer
bash build_apptainer.sh
```

To use the container for training run the following command:

```
apptainer run --bind /data:/data --nv \
  /work/gnndebugger/apptainer/dbgc.sif \
  --config-path /data/tconfig \
  --config-name train.yaml paths.data_dir=/data
```

For the LJA with ML inference

```
apptainer run --bind /data:/data
/work/LJA/apptainer/dbgc.sif
    --reads reads.fastq \
    -o /data/assemblies --threads 32 \
    --CoverageBasedCorrection.reliability-mode multiplicity \
```

```
--CoverageBasedCorrection.ml-threshold 0.85 \
--TopologyBasedCorrection.reliability-mode multiplicity \
--TopologyBasedCorrection.ml-threshold 0.7
```
